# Tiger diet in Ranthambore Tiger Reserve: how do metabarcoding and mechanical sorting compare?

**DOI:** 10.1101/2025.06.18.660266

**Authors:** Sarang Mishrikotkar, Nidhi Yadav, Abishek Harihar, Varun R. Goswami, Uma Ramakrishnan, Mousumi Ghosh-Harihar

## Abstract

Accurately describing large carnivore diets is critical for understanding trophic interactions and identifying targeted conservation strategies. Most studies have relied on traditional dietary analysis based on mechanical sorting and identification of undigested prey remains, a method known to be error-prone and ecologically biased. Here, we compare the diet of tigers using non-invasively collected scats from Ranthambore Tiger Reserve in India, analysed using mechanical sorting and DNA metabarcoding. We found that DNA metabarcoding outperformed mechanical sorting in detecting higher overall prey and rare prey species occurrences, uncovering higher prey diversity. The study revealed that tigers were subsisting mainly on wild prey such as sambar and chital. However, domestic cattle contributed the highest relative prey biomass to their diet. Our findings demonstrate that DNA metabarcoding is an efficient, effective and powerful approach that overcomes several of the previously identified biases of the mechanical sorting approach and provides an accessible and particularly useful tool for carnivore dietary studies based on non-invasive samples. The increased frequency of livestock depredation relative to previous studies highlights the need for active mitigation measures to secure this population.

## Introduction

Large carnivores, as apex predators, disproportionately impact ecosystems and food webs through prey limitation and top-down trophic cascades (Paine, 1980; Ripple et al., 2014; Atkins et al., 2019; Ritchie et al., 2012). Representing some of the most imperilled species worldwide owing to their dietary specialisation and wide-ranging habits, their numbers strongly influence prey species’ density and biomass (Cardillo, 2003; Ripple et al., 2014; Hatton et al., 2015). Therefore, accurately assessing their diet is an essential first step in understanding their role in ecosystem processes (e.g., resource utilisation, trophic interactions, predator-prey dynamics, interspecific niche partitioning) and their life-history strategies (Miquelle et al., 1996; Soulé et al., 2003; Sheppard & Harwood, 2005; Estes et al., 2011; Wallach et al., 2015; de Sousa, Silva, & Xavier, 2019). Furthermore, rigorous dietary analyses are also critical from an applied perspective since livestock depredation and prey depletion are known to be key drivers of the global decline in and local extinction of carnivores (Ceballos & Ehrlich, 2002; Treves & Karanth, 2003; Wolf & Ripple, 2016).

Since observational studies are rather challenging due to most carnivores being rare and elusive (Long et al., 2007), diet analyses traditionally relied on mechanical sorting and morphological and microhistological identification of undigested prey remains (e.g., hair, feathers, exoskeleton, and bones) in non-invasively collected scat samples (Spaulding, Krausman, & Ballard, 2000; Henschel, Abernethy, & White, 2005; Andheria, Karanth, & Kumar, 2007; Klare, Kamler, & Macdonald, 2011). However, this approach has several limitations, including being time and labour-intensive, and requiring strong taxonomic skills and prior reference keys (Reynolds & Aebischer, 1991; Lake, Burton, & van den Hoff, 2003; Sheppard & Harwood, 2005; Packer et al., 2009). Relying on the discovery of undigestible body parts, rare species and species with soft parts are often missed or misidentified (Spaulding, Krausman, & Ballard, 2000). Similarly, misidentification occurs for some species that are difficult to distinguish based on hard parts (Zeale et al., 2011). Observer bias also leads to erroneous ecological inferences (Spaulding, Krausman, & Ballard, 2000; Massey et al., 2021). Finally, field identification of predator scats without molecular validation is another source of erroneous and biased inferences in dietary studies (Morin et al., 2016; Weiskopf, Kachel, & McCarthy, 2016).

In recent years, metabarcoding of fecal DNA has emerged as a reliable alternative for dietary analysis ((Shehzad et al., 2012; De Barba et al., 2014; McInnes et al., 2017; Monterroso et al., 2019; Thuo, 2020; Ghosh-Harihar et al., 2024). DNA metabarcoding combines high-throughput sequencing (HTS) with “universal markers” that target short mitochondrial DNA fragments to maximize detection from the widest possible range of prey species, allowing the sequencing of millions of DNA fragments from multiple species simultaneously (Valentini, Pompanon, & Taberlet, 2009; Razgour et al., 2011; Taberlet et al., 2012; Soininen et al., 2015). Furthermore, the use of novel bioinformatics tools for the selection of appropriate molecular barcodes, data curation, and robust HTS filtering allow finer taxonomic resolution of dietary components from fecal DNA, enabling a more accurate description of diet composition (Valentini, Pompanon, & Taberlet, 2009; Brown et al., 2012; Corse et al., 2017). Comparisons between mechanical sorting (i.e., morphological analysis of undigested remains) and metabarcoding suggest that generally, metabarcoding reveals a higher prey diversity, detects rarer species often overlooked by mechanical sorting, and is more effective for even degraded scats (Berry et al., 2015; Massey et al., 2019)-though some suggest a more complementary role for these approaches in dietary investigations (Pertoldi et al., 2021).

Dietary baselines are critically needed to conserve wild tigers to understand their foraging ecology, responses to anthropogenic changes in habitat and fragmentation, assessment of recovery potentials, and understanding the effects of declining prey and the contribution of livestock to diets. Tigers prefer ungulate prey and show selective size-based predation, with the preferred weight range of prey being 60 to 250 kg (Hayward, Jękedrzejewski, & Jêdrzejewska, 2012). Their diet, particularly differential prey selection in terms of species, body weight, sex, and age, also plays a crucial role in facilitating coexistence with other sympatric predators such as the common leopard Panthera pardus and dhole Cuon alpinus (Karanth & Sunquist, 1995; Andheria, Karanth, & Kumar, 2007; Harihar, Pandav, & Goyal, 2011; Lahkar et al., 2021a). Furthermore, demographic responses to human-induced prey depletion or size-selective prey poaching are mediated through changes in diet (Ramakrishnan, Coss, & Pelkey, 1999; Linkie & Ridout, 2011; Miller, Ament, & Schmitz, 2014; Sugimoto et al., 2016; Karanth et al., 2017; Steinmetz et al., 2021). These insights were gained from dietary studies primarily based on morphological and microscopic examinations of scats and kills through mechanical sorting of undigested remains. However, given a more recent recognition that such methods might not accurately reflect carnivore diets, investigating how a metabarcoding approach can improve our understanding of diet is crucial.

Here, we focused on assessing the diet of wild tigers in Ranthambore Tiger Reserve in India while providing a formal comparison of diet analysis using mechanical sorting and DNA metabarcoding. We opportunistically collected scats across the tiger reserve using pre-existing routes that were sampled repeatedly. By analysing the scats processed using both approaches, we assess if metabarcoding uncovered a higher prey diversity, resulted in finer taxonomic resolution, and could identify rarely consumed prey. Finally, we describe the dietary profile of these tigers based on findings from DNA metabarcoding.

### Methodology Study area

We conducted this study in the Ranthambore Tiger Reserve (RTR), Rajasthan, India. The vegetation of this region is tropical dry deciduous forest and tropical thorn forest (Champion & Seth, 1968), with dhok (Anogeissus pendula) as one of the dominant tree species spread across the landscape. RTR is the only source population of wild tigers in the Western Indian Landscape (Ranthambore-Mukundara-Kuno-Shivpuri-Madhav), with a density of ∼ 7.5 individuals / 100km^2^ (Sadhu et al., 2017). Tigers in RTR are genetically isolated from other landscapes in the country (Natesh et al., 2019). Prey species occurring within the study area include chital (Axis axis), sambar (Rusa unicolor), nilgai (Boselaphus tragocamelus), langur (Semnopithecus entellus), and wildpig (Sus scrofa), and chinkara (Gazella bennetti). Dietary analyses conducted two decades ago using mechanical sorting revealed a preference for large prey (107–114 kg), with chital and sambar being the preferred prey of tigers (Bagchi et al. 2003). The study also found that nilgai and chinkara were consumed less than their availability, and livestock comprised 10–12 % of their diet.

### Sample collection

We opportunistically collected tiger scats along the forest road network in five planned collection surveys between December 1^st^ 2019 and February 15^th^ 2020, to collect multiple replicates of tiger scats (Fig. 1). On encountering a scat during the survey, we first estimated the age (fresh -wet from outside or inside; old-completely dry) based on appearance (Ciucci et al. 1996). We collected two different types of samples for dietary analyses from each scat assigned as fresh. For diet metabarcoding, we collected multiple biological replicates (4–5) from the cross-sectional area of each scat to minimize tiger DNA (Stenglein et al. 2010, Massey et al. 2019). We stored each sample in an individually labeled and small zip lock bag, bagged in a larger zip lock bag with 2–3 silica gel crystals. All the genetic samples were stored at - 21°C in the field and transferred back to the National Centre for Biological Sciences, Bengaluru, on dry ice for storage at -21°C until DNA extraction.

**Figure 1:**
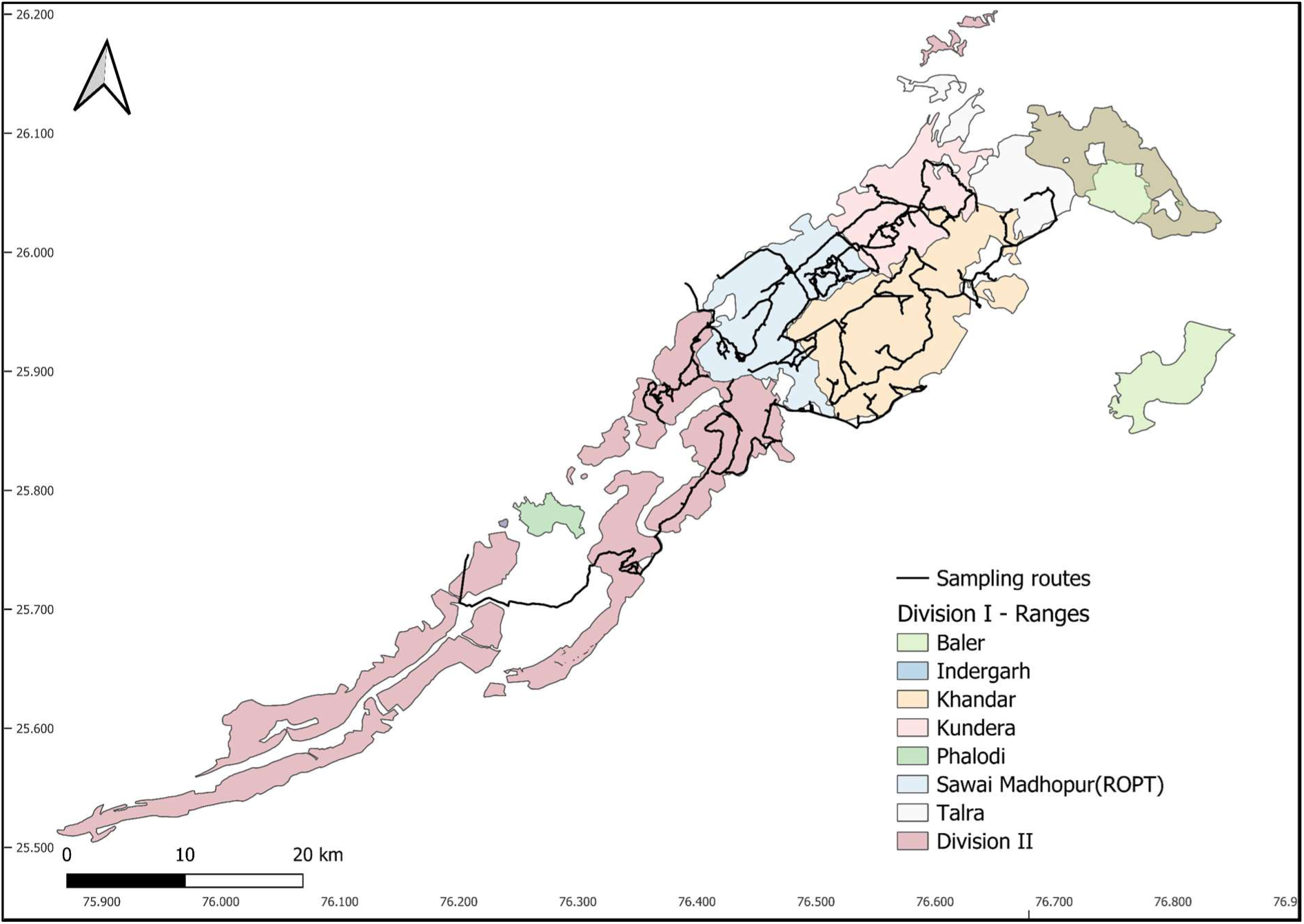
Sampling routes used for collecting tiger scat samples within the Ranthambore Tiger Reserve, India. The names indicate the forest ranges in RTR.

A portion of the scat with such undigested remains was collected and stored in a large airtight bag for dietary analysis based on mechanical sorting and identifying undigested prey remains. We also sampled the epithelial cell tissue from the scat surface to confirm the predator identity using a sterile swab dipped in Longmire lysis buffer (Longmire et al., 1997). We then broke the tip of the swab and collected it in a 2 ml vial containing Longmire buffer. To minimize the risk of contamination with human DNA, we wore sterile disposable gloves while collecting these samples.

### Mechanical sorting of prey remains

We first separated undigested prey remains by washing the collected scat samples under running water through a fine (< 1mm) metal wire sieve. The material left in the sieve, which included undigested hair particles, hooves, and other organic material, were then dried in the sun on paper for three days to avoid fungal and bacterial growth. We randomly selected 20 prey hair strands from each dried sample, mounted them in xylene on a glass slide with a coverslip, and examined them under a microscope (LABOMED Lx400 - 10*40X & 10*100X magnification). We examined the observed medullary pattern of the hair to compare them to a reference library of published images of species-specific medullary patterns to identify the prey species (Bahuguna, 2010; Bhawan et al., 2016; Upadhyaya et al., 2018).

### Predator identification

We extracted DNA from the swab samples using a DNA blood and tissue kit (QIAGEN, Germany) following manufacturers’ instructions in a dedicated facility for extracting DNA from non-invasive samples. To identify the host species, we amplified a 202bp of the 16S rRNA region of the mitochondrial DNA using a felid-specific primer (Mukherjee et al. 2016). The sequences are as follows: 16S rRNA-F (5’-AATTGACCTTCCCGTGAAGA-3’), 16SrRNA-R (5’-TCCGACTGGTTAGTCTAGAT-3’). The optimum PCR conditions included an initial denaturation at 95° C for 15 min, followed by 45 cycles of denaturation at 94° C for 30 sec, annealing at 57° C for 30 sec, extension at 72° C for 30 s, and a final extension at 72° C for 10 min, hold at 10° C. The PCR products were then electrophoresed on a 2% agarose gel, and the samples with successful amplification were sequenced using Sanger sequencing. We performed additional BLAST hits in GENBANK with a minimum query coverage of 98% to confirm the identification of tiger-positive samples, which were further used for determining diet.

### Diet data generation

Since scat samples contained different components of undigested prey like hair, bones, hooves, etc., we homogenized each scat sample into a uniform powder using mortar and pestle after freezing the sample with the help of liquid nitrogen. It was then immediately transferred to a 2 ml centrifuge tube/vial for overnight digestion at 56°C. For DNA extraction, we followed the manufacturer’s instructions using the QIAamp® Fast DNA Stool Mini Kit (QIAGEN, Germany) with a slight modification (InhibitEX Buffer) and eluted the DNA in a total volume of 100 µl. Standard precautions and controls were included to avoid and detect contamination. We prepared the library for diet metabarcoding following the steps described in Ghosh-Harihar et al. (2021) with slight modifications (see Supplementary information). First, we amplified each sample using the metabarcode 12SV5 (Riaz et al. 2011), adding a blocking oligonucleotide PantB to repress the amplification of dominant host tiger DNA (Shehzad et al., 2015). We modified all the primers with the addition of the Illumina overhang adapters. Therefore, we also amplified several negative (extraction and PCR) and positive (mock prey community sample) controls along with the samples. The positive control was a mock prey-predator community sample containing DNA from gaur (Bos gaurus), spotted deer (Axis axis), and dog (Canis sp.) in equal concentrations along with the predator DNA (tiger Panthera tigris) at four times the concentration. We sequenced the final libraries (including indexing controls) using the Illumina MiSeq platform (Illumina, USA). We analyzed and filtered the HTS data following Ghosh-Harihar et al. (2021) (see Supplementary information).

Finally, we employed several stringent steps to confirm the assignment of sequences to unique molecular taxonomic units (MOTU). We used BLAST on NCBI separately for sequences not identified up to the species levels (E-value threshold: 1e-10; minimum query coverage: 98%). Additionally, we relied on biogeographic knowledge of the prey community in the study area to re-assign a taxonomic identity to closely related species.

### Diet composition

Prey composition in tiger diet was described using the frequency of occurrence (FOO) and relative biomass (B). Frequency of occurrence (FOO) was calculated as the proportion of scats in which the species was detected, as prey remains in mechanical sorting method or sequence reads in metabarcoding. Different prey species’ contribution to the tiger diet was determined using relative biomass. This was calculated using a generalized model based on an asymptotic, allometric relationship: biomass consumed per collectible scat (Y) for each prey species = 0.033–0.025exp^-4.284(prey^ ^weight/predator^ ^weight)*^predator weight, applicable to all obligate carnivores to compute prey biomass consumed from scats (Chakrabarti et al., 2016). Using the correction factor, we computed the relative biomass (B) with the following equation:

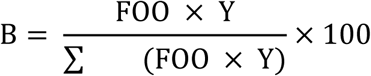

The predator and prey weights used in this calculation were derived from multiple studies (Biswas & Sankar, 2002; Carbone & Gittleman, 2002; Bagchi, Goyal & Sankar, 2003; Cordier et al., 2012; Lahkar et al., 2021; Endo et al., 2022) (Table S1).

### Comparative analysis

Discrepancies in the output from two methods, metabarcoding and mechanical sorting, were analyzed based on the number of prey species detected in overall scat samples. The average number of prey species encountered per scat was compared across methods using a non-parametric Wilcoxon rank-sum test with continuity correction. To assess which prey species contributed the most to the observed dissimilarity between the output from the two methods, we used similarity percentage analysis (SIMPER). A subset of scats analyzed with both methods (n=156) were compared by determining FOO and relative biomass. Finally, we present the frequency of occurrence and relative biomass from the metabarcoding analysis to describe the diet of tigers in RTR.

## Results

We surveyed the landscape five times using the road network, covering a total distance of 2,220 km, and collected 207 unique scat samples, each collected in replicates (total n = 414 subsamples). Of the 207 samples collected, 193 (93.2%) were confirmed as tiger scats, 11 (5.3%) as leopard scats, and 3 (1.5%) could not be assigned to as any specific predator. We finally used data from 156 scat samples that were analyzed using diet metabarcoding and mechanical sorting. The variation in the frequency of occurrence for each species stabilized around 50 scat samples for mechanical sorting and metabarcoding (Fig. S1). Therefore, the sample size of 156 was deemed sufficient to assess the diet of the tiger population in RTR.

### Diet metabarcoding versus mechanical sorting

We compared the diet of tigers from the 156 scat samples that were analyzed with both mechanical sorting and metabarcoding.

In metabarcoding analysis, post High Throughput Sequencing (HTS) filtering, these 156 samples retained a total of 9,060,458 sequences for dietary analysis (Table S2). We used PantB blocking primer along with the universal vertebrate primer 12SV5, and found that **19.6%** of the sequences belonged to the predator species. Comparatively, metabarcoding detected 225 more prey occurrences and higher number of unique Molecular operational taxonomic units (mOTUs) across the 156 scat samples (Fig. 2a).

**Figure 2:**
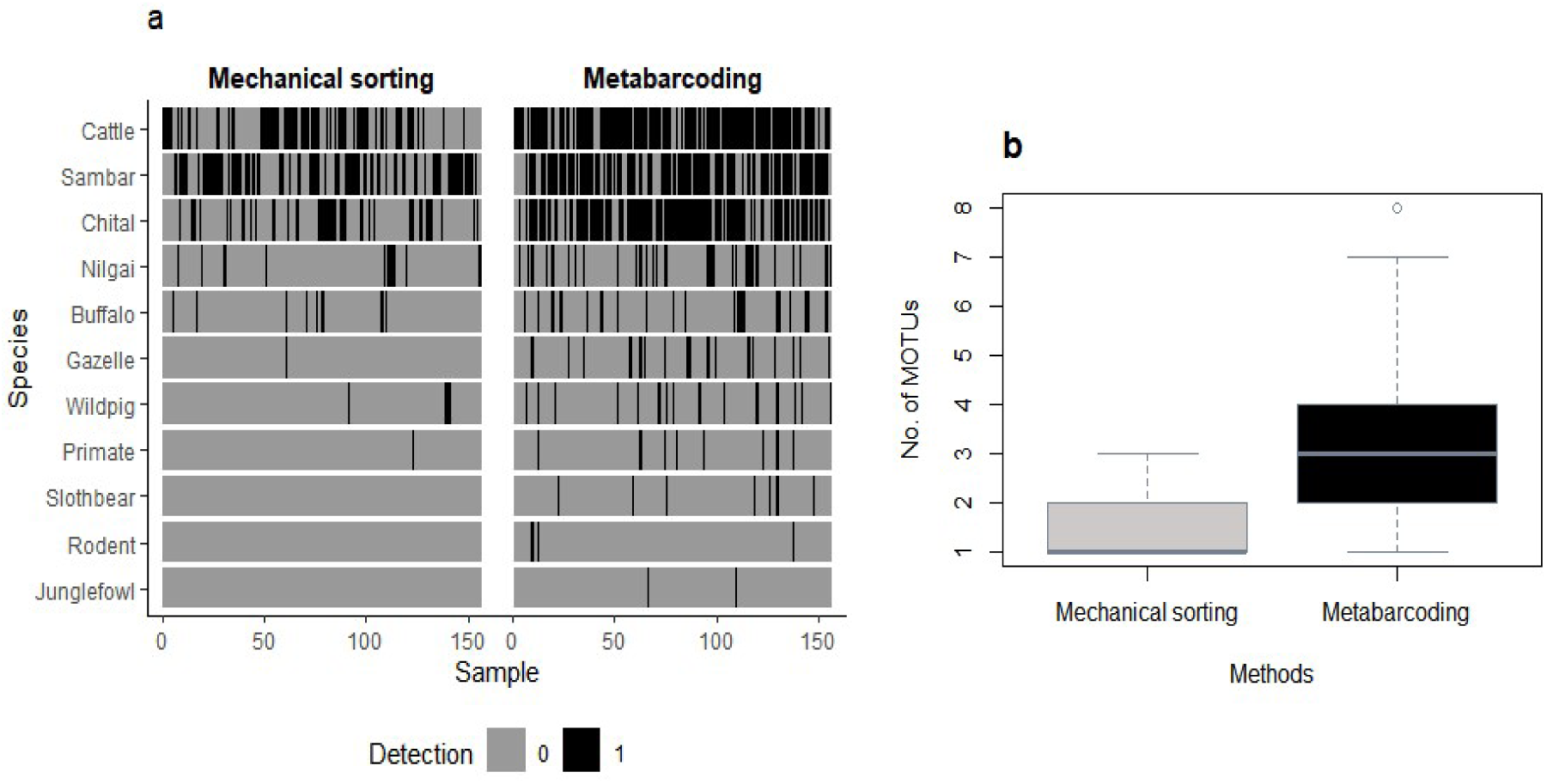
(a) Detections of prey species MOTUs using mechanical sorting and metabarcoding. (b) The number of MOTUs detected per tiger scat sample using the two methods of dietary analyses.

The mean number of species per sample was also higher for metabarcoding than mechanical sorting (Wilcoxon signed-rank test, V = 114, p-value < 2.2e-16, Fig. 2b).

On average, each sample contained 1.32 (± 0.56 SD) prey species per scat, a maximum of three species and a total of eight unique prey species when using mechanical sorting (Fig. 2b). Metabarcoding detected eleven unique prey taxa, a maximum of seven diverse prey in a sample, and an average of 2.93 (± 1.36 SD) prey species per scat, with most of the scats (65.4%) containing two–three prey species (Fig. 2b). Metabarcoding also revealed three rare species that were undetected during mechanical sorting, including sloth bear (Melursus ursinus), a rodent species and red jungle fowl Gallus gallus. There were no instances of mechanical sorting identifying a prey species, which was not detected by metabarcoding.

The frequency of occurrence (FOO) figures of different prey taxa was relatively similar. However, absolute FOO values were much higher for each taxon when using metabarcoding (Fig. 3a). Additionally, the order of the most frequently detected species differed between the two methods. Metabarcoding detected cattle most frequently (76.28%), followed by chital (68.6%) and sambar (64.1%). On the other hand, sambar appeared to be the most frequently detected species (50.14%), followed by cattle (41.66%) and chital (23.71%) when using mechanical sorting. The SIMPER analysis also indicated that these three species contributed most (72.4%) to the observed differences between the two approaches (Table S3).

**Figure 3:**
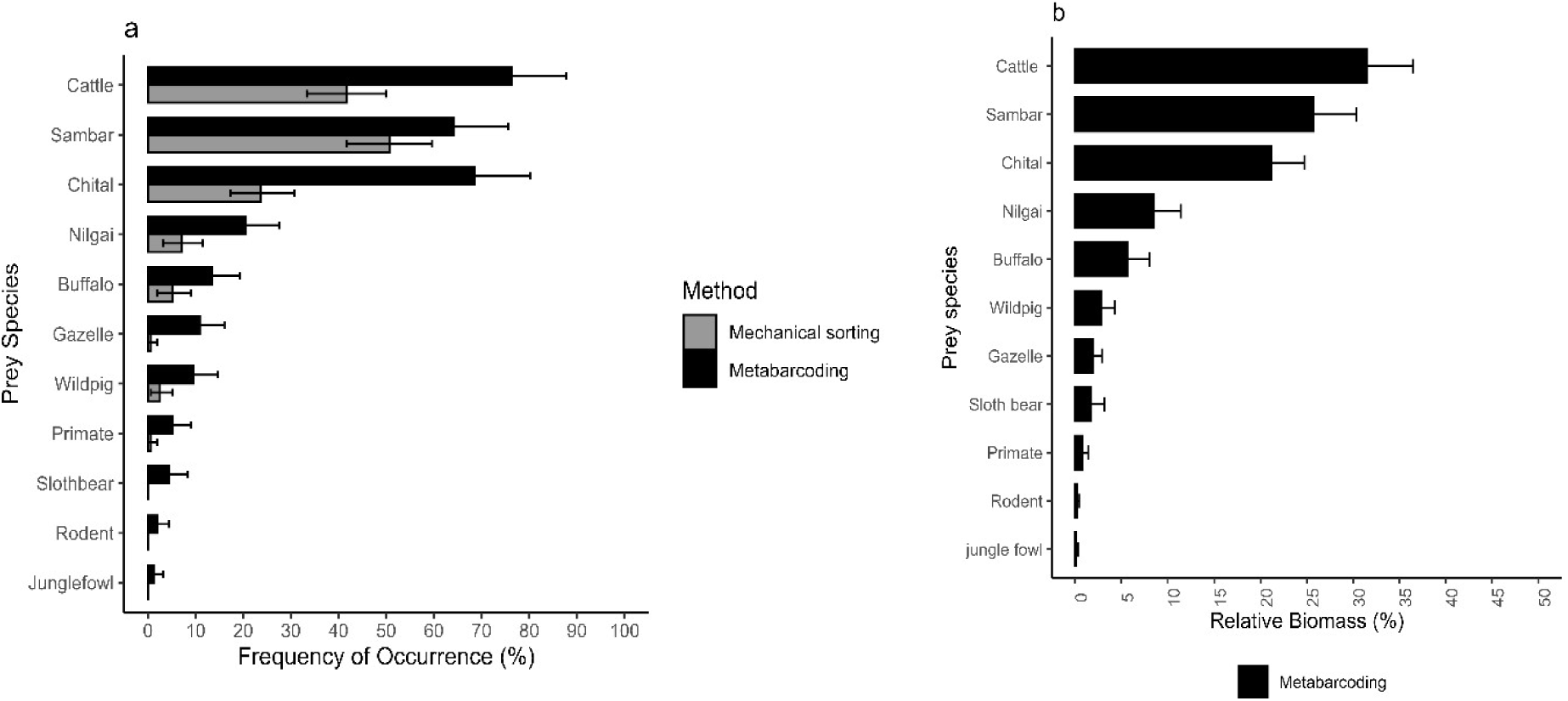
(a) Frequency of occurrence (FOO) of prey taxa in tiger diet across metabarcoding and mechanical sorting. (b) Relative biomass of prey species in tiger diet using metabarcoding.

### Diet of tigers based on metabarcoding

We detected 431 prey occurrences and identified 11 unique prey taxa using diet metabarcoding. Domestic cattle was the most frequently occurring prey species detected in 76.28% of all the scat samples and comprised 32.13% of the diet by biomass. Domestic water buffalo was also detected in ∼14% of the scats. Among the wild prey species, chital was detected most frequently, followed by sambar, nilgai, wild pig, chinkara, and primates (Fig. 3a). The other rarer prey species, including sloth bear, rodent, and jungle fowl, were detected in <5% of the scats. Out of the eleven prey species detected using metabarcoding, three species (cattle, sambar, and chital) contributed most (>78%) towards the relative prey biomass consumed (Fig. 3b). Wild prey contributed nearly 61.5% to the tiger’s diet in relative biomass consumed, while domestic species (cattle, buffalo) contributed the remaining 38.5%.

## Discussion

In our study, DNA metabarcoding outperformed mechanical sorting in uncovering higher prey diversity and overall prey and rare prey species occurrence. Our results suggest that tigers in RTR largely subsist on wild prey, although domestic cattle contribute the highest relative prey biomass to their diet.

### Methodological insights

The methodological comparisons between DNA metabarcoding and mechanical sorting largely highlight the biases associated with the traditional mechanical sorting protocol followed for studying tiger diet. For instance, mechanical sorting is unlikely to detect very small-bodied prey species and any species that is consumed as soft tissue (e.g. in case of scavenging). In case of rare species or hair that undergo morphological alterations after passing through the digestive tract, observers tend to assign them to the most commonly occurring prey species (Shehzad et al. 2012; Shores et al. 2015). This leads to overestimation of certain common prey species in the diet (Massey et al. 2019). Rare species are more likely to be overlooked since only a few randomly undigested remains (mostly hair) are selected for microscopic examination in most studies. Furthermore, manual comparison of microscopic images with reference images can potentially introduce observer bias due to fatigue and lead to erroneous identification. In our case, we did not include sloth bear and jungle fowl in the reference database for mechanical sorting, and hence, missed detecting these species that occurred very rarely (<4% of the samples). Together, these factors might explain the fewer prey occurrences as well as lower number of prey species detected using mechanical sorting. This has been demonstrated in the case of wolf scats, where both the diversity of prey species and occurrences were higher when using diet metabarcoding (Berry et al. 2015; Massey et al. 2021). However, unlike other such comparisons, we did not find evidence for false positives in case of mechanical sorting (Massey et al. 2015) or complementarity in detections between the two approaches (Pertoldi et al. 2021).

Tiger diet has been studied primarily using mechanical sorting of undigested prey remains in non-invasively collected samples (Bagchi et al. 2003; Andheria et al. 2007; Harihar et al. 2011; Lahkar et al. 2021) or examination of kills (Miller et al. 2014). Our findings suggest that diet metabarcoding is the more powerful approach to assessing tiger diets from non-invasively collected scat samples. So far, only one study has employed cytochrome oxidase subunit-1 or COI DNA metabarcoding to describe the prey preferences of the Malayan tigers although the sample size was rather small and taxonomic resolution was poor (Gani et al. 2024). Use of a COI metabarcode yielded just 3.56% of chordate sequences unlike our study where we used a vertebrate-specific primer in combination with a blocking oligonucleotide to suppress predator sequence amplification, which yielded much better taxonomic resolution. Furthermore, we found that nearly 5% putative tiger scats were identified as leopards using molecular identification highlighting the importance of confirming predator identification using molecular markers and not relying on morphological identification alone.

While DNA metabarcoding clearly outperforms mechanical sorting in detecting prey diversity based on presence–absence data, estimating relative biomass based on read counts remains a methodological challenge. In this study, we adopted a conservative approach by calculating relative biomass using the frequency of occurrence (FOO) rather than sequence read proportions since the read counts did not reflect the DNA concentrations in our mock community samples. This is a well-recognised limitation of diet metabarcoding studies, owing to factors such as differential digestion, gene copy number variation, and PCR biases (Deagle et al., 2019). Converting read counts reliably into relative biomass consumed would require more careful evaluations using controlled feeding trials on captive individuals.

### Diet of tigers in RTR and implications

Large herbivores including chital, sambar, domestic cattle, nilgai and buffalo formed the bulk of the tiger diet contributing > 90% of the relative biomass (Fig. 3b), which is in concordance with previous studies (Karanth & Sunquist 1995; Bagchi et al. 2003; Hayward et al. 2012; Andheria et al. 2007; Harihar et al. 2011). In our study the two large cervids (chital and sambar) contributed 44.6% to the diet of tigers in RTR, which is lower than the relative biomass contributed by these species (78%) reported in a previous study in the area (Bagchi et al. 2003). Domestic cattle and buffalo comprised 37% of the relative biomass consumed by tigers in our study, compared to just 11.08% reported in Bagchi et al. (2003). Though not strictly comparable due to the previous study being situated in a much smaller area within the National Park and using mechanical sorting, an increase in livestock in the diet of tigers in RTR appears plausible based on these findings. But irrespective of a temporal trend, our results seem to suggest that livestock make up a considerable portion of the diet of tigers in RTR today.

The depredation of domestic livestock (cattle, buffaloes, goats) has been recognized as a significant part (88.5%) of human–tiger conflict in the RTR Landscape (Singh et al. 2015). Data collected by the forest department suggests that cattle were attacked most often (69.4%), followed by buffaloes (19.3%) and goats (11.4%). Our samples also show a higher frequency of occurrence of cattle in the diet of tigers compared to buffaloes. We did not encounter goats in the diet during our sampling duration. A vast majority of these incidents were recorded in either inside the villages or in the agricultural fields peripheral to the park boundary, suggesting that we should expect tiger scats containing livestock to be concentrated along the park boundary. However, in our study scats containing livestock were not confined to the periphery, but were distributed throughout the park (Fig. 4). This could be potentially attributed to the steady increase in tiger density ((Sadhu et al. 2017) and the mostly linear shape of the park, which could be facilitating the movement of many individual tigers into the surrounding villages. Furthermore, we occasionally encountered free ranging cattle grazing inside the park while sampling, which likely contributed to the presence of livestock in diet in the core area. This poses a significant challenge to conserving tigers in this localized, remnant, semi-arid population in western India. Conflict mitigation in the case of isolated and mostly linear parks like RTR hinges on mechanisms that can negate losses and foster coexistence, including measures such as adequately compensating affected villagers, improved approaches to guarding livestock, and potentially setting up insurance schemes (Goodrich et al. 2010; Singh et al. 2015). Adopting a participatory approach with the local communities to identify suitable conflict mitigation strategies and robust monitoring of the effectiveness of mitigation strategies would be crucial to the long-term success of such measures (Treves et al. 2009; Goodrich et al. 2010).

**Figure 4:**
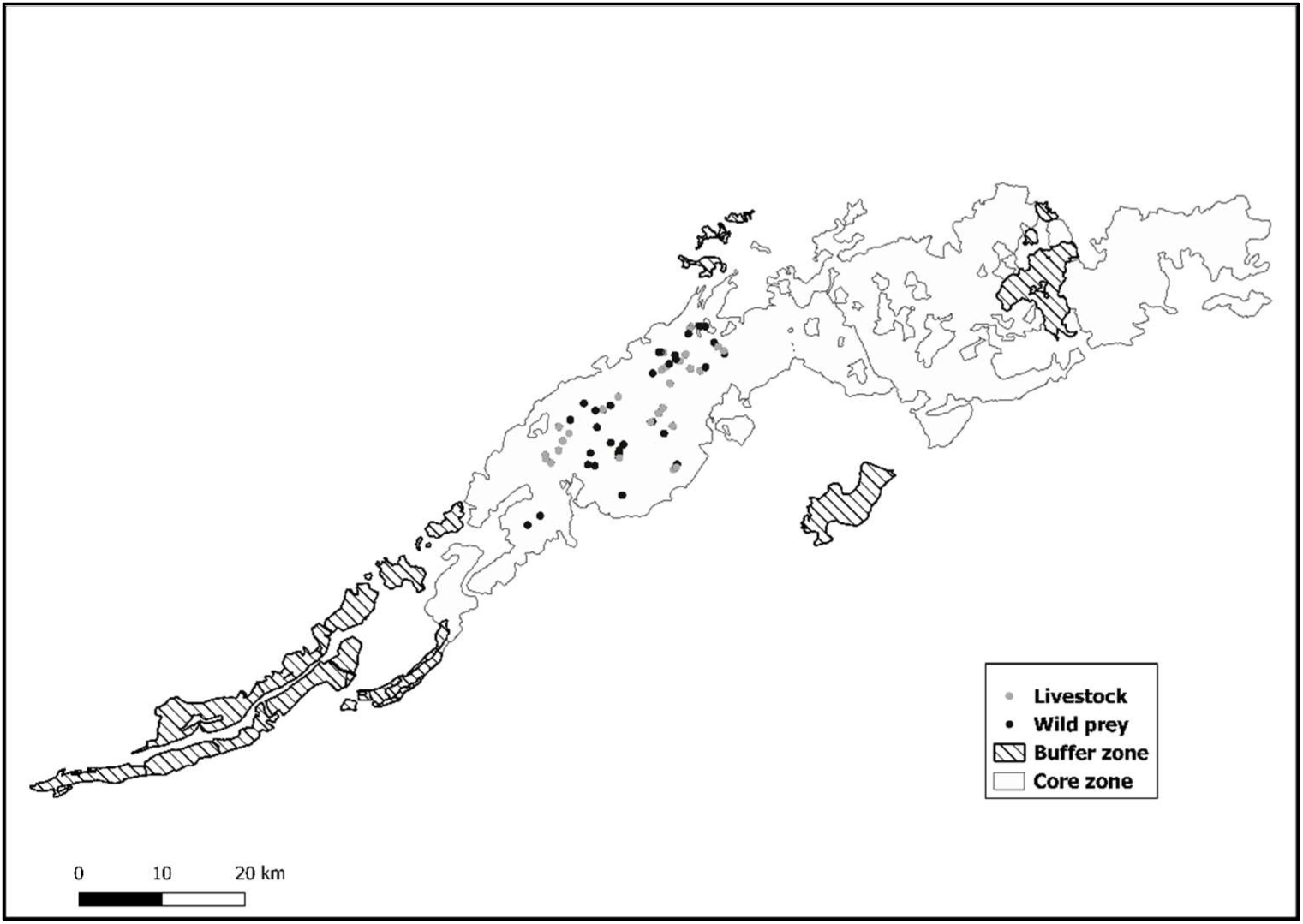
The location of scat samples containing livestock (with and without wild prey) and those with only wild prey remains across RTR.

## Conclusions

Our results demonstrate that DNA metabarcoding is a more efficient, effective and powerful approach to characterize the diet of tigers from non-invasively collected scat samples when compared to the traditional mechanical sorting approach. Diet analysis remains a key tool to understand prey-predator interactions, species interactions and species response to changes in the environment. In a changing world, under the dual influence of rapid human development and global climate change, identifying tools that are effective, sensitive and are less error-prone is critical in our ability to monitor changes in species responses and interactions. DNA metabarcoding is clearly one such method that overcomes many of the previously identified biases of mechanical sorting and provides an accessible and highly effective tool for carnivore dietary studies based on non-invasive samples. Through this approach, we were able to reveal a higher frequency of livestock depredation among tigers in RTR than previously reported, a finding of high conservation value that calls for participatory and long-term conflict mitigation to secure an important but vulnerable population of an endangered large carnivore.

## Acknowledgements

We are grateful to the National Centre for Biological Sciences–Tata Institute of Fundamental Research and the Department of Atomic Energy for institutional, financial, and logistical support. We sincerely thank the Rajasthan Forest Department for granting research permissions and providing logistical assistance during the study. Special thanks to Mujahid Khan, Krishna Prajapat, Mujeeb Ahmad, and interns Tejaswini J. and Vijay Naidu for their dedicated support in the field. We also extend our gratitude to our lab mates—Vinay Sagar, Anubhab Khan, Megan Aylward, Vanessa Paynter, Aditi Prasad, Ryan Rodrigues, Abhinav Tyagi, Rupsy Khurana, Anuradha C., and Mayuresh Gangal—for their invaluable inputs.

## Author Contributions Statement

SM, MGH and UR conceived the ideas and designed methodology; SM and NY collected and processed the data; SM, NY, AH, MGH and VRG analysed the data; MGH and SM led the writing of the manuscript. All authors contributed critically to the drafts and gave final approval for publication.

