## Supplementary material for "Tiger diet in Ranthambore Tiger Reserve: how do metabarcoding and mechanical sorting compare?"

**Supplementary Information**Table S1: Tiger and prey species body weights used in calculating relative biomass consumed following Chakraborti et al. (2016).

| **Prey species** | **Body mass (kg)** | **Source** |
| --- | --- | --- |
| Tiger | 180 | (Carbone and Gittleman, 2002) |
| Sambar | 125 | (Bagchi et al., 2003) |
| Cattle | 180 | (Bagchi et al., 2003) |
| Chital | 45 | (Bagchi et al., 2003) |
| Wild pig | 38 | (Bagchi et al., 2003) |
| Sloth bear | 90 | (Biswas and Sankar, 2002) |
| Nilgai | 180 | (Bagchi et al., 2003) |
| Buffalo | 273 | (Bagchi et al., 2003) |
| Jungle fowl | 0.985 | (Endo et al., 2022) |
| Chinkara | 12 | (Bagchi et al., 2003) |
| Rodent | 0.5 | (Lahkar et al., 2021) |

a. **Library preparation and sequencing for diet metabarcoding**

We prepared the library for diet metabarcoding in three steps that consisted of two PCR reactions and a pooling step using MiSeq Reagent Kit v2, DNA library preparation kit. Following DNA extraction, each sample was amplified using primer pair 12SV5F, 12SV5R (Riaz et al., 2011)*.* The primer targets approximately 100 base pairs of the V5 loop in the 12S region of the mitochondrial rRNA gene common in all vertebrates. To repress the amplification of dominant host tiger DNA, we also added a blocking oligonucleotide, PantB (Shehzad et al., 2015). The reaction was performed in a total volume of 25 ul and consisted of 12.5ul of 2X master mix (Qiagen Multiplex PCR plus kit), 2.5ul of 250nM primer pool, and 4 ul of template DNA. We included negative control in the last well for each row to detect any contamination due to pipetting error. We also added 2 positive controls in each plate to check the efficiency of primer pairs and blocking oligonucleotide. The positive control was a mock prey community; Gaur (*Bos gaurus*), spotted deer (*Axis axis*), Dog (*Canis sp.*), all in equal concentrations along with the predator DNA; tiger (*Panthera tigris*) at four times the concentration to that of the prey community. PCR conditions were set on an initial denaturation step at 95° C for 15 min, followed by 5 cycles of denaturation at 95° C for 30 sec, annealing at 57° C for 30 sec, extension at 72° C for 2 min with 5% ramp and 40 cycles of denaturation at 95° C of 30 sec, annealing at 65°C for 60 sec, and extension at 72° C for 30 sec. To check the amplification in each sample, gel electrophoresis was performed on 2% agarose gel and the overall DNA concentration was also quantified using Qubit Fluorometer (Invitrogen, Darmstadt, Germany).

To remove the unused primer and primer dimers, PCR amplicons were purified using 37.5 µl of re-suspended Agent AmPure XP Beads (Beckman Coulter, Fullerton, CA, USA). To tag the samples for multiplexing, we added dual indices and Illumina overhang adapters (forward overhang: 5’ TCGTCGGCAGCGTCAGATGTGTATAAGAGACAG-primer sequence; reverse overhang: 5’ GTCTCGTGGGCTCGGAGATGTGTATAAGAGACAG‐primer sequence) from Nextera XT Indexing Kit. The limited cycle PCR was carried out in a volume of 50 µl with 25 µl of Qiagen Multiplex Kit Master Mix (Qiagen) and 5 µl of each indexing primer (i5 and i7) along with 20ng of the purified template DNA. The PCR conditions included an initial denaturation step at 95 ° C for 3 min, followed by 8 cycles of denaturation at 95 ° C for 30 s, annealing at 55 ° C for 30 s, and extension at 72 ° C for 30 s, followed by a final extension step at 72 ° C for 5 min. At the end of this step, each sample was assigned a unique combination of 12 i5 and eight i7 index that enabled multiplexing 96 different samples in single Miseq run. This was followed by a second purification step in which 50 µl of DNA template was used with 50 µl Agencourt AMPure XP beads (Beckman Coulter, Fullerton, CA, USA)(1.5x). To check the final fragment size of the recent PCR product and to verify any potential primer contamination, 1 µl (1:50 dilution) of the dual-indexed library was analyzed on a 2100 Bioanalyzer and Agilent High Sensitivity DNA chip (Agilent Technologies, Santa Clara, USA).

b. **Miseq Sequencing**

Miseq platform (Schirmer et al., 2016) was used for sequencing the library after pooling it by 5µM per sample. NaOH, diluted with a hybridization buffer, was used to denature the library. 6pM of this along with 10% PHiX control was loaded in a Miseq flow cell with a 300-cycle Reagent Kit v2 (Illumina). The paired-end sequencing was run at 2x150bp to yield high-quality reads in pairs for each sample.

c. **HTS filtering**

We analysed the raw sequence reads with a standardized bioinformatics pipeline, OBITools (Boyer et al., 2016)*.* The raw reads for each sample were paired using *Illuminapairedend*. For assigning each sequence to corresponding primer and index, *ngsfilter* was used (Shehzad, 2012). Sequences with counts lower than 10 and shorter than 80 bps were discarded. We filtered the sequencing errors or variants during PCR using *obiclean*. Based on a directed acyclic graph (Ficetola et al., 2015), sequence variants, classified as “internal” with a threshold of 5% were removed from the dataset.

Sequences were assigned taxonomic identity using BLAST against 12S vertebrate sequences in a reference database ( Ghosh-Harihar et al. 2019)*.* The reference database was assembled using the *ecoPCR* program (Ficetola et al., 2015) and built using the sequences derived from the European Molecular Biology Laboratory (EMBL) database. We used *ecotag* to assign a taxonomic identity to each sequence that corresponded to the last common ancestral node in the NCBI taxonomic tree of sequences from the reference database.

A series of filtering steps using multiple Low-frequency noise (*LFN)* threshold was carried out on the taxonomically assigned sequences to eliminate false-positives due to contamination, mistagging and PCR/sequencing errors (De Barba *et al.*, 2014; Corse *et al.*, 2017). To increase the accuracy in the taxonomic assignment dataset, sequences that mapped on to the reference database with a threshold of more than 95% identity were retained.

LFN_neg_: A threshold calculated using the most frequent read count in the negative controls was used to filter the dataset and any sequence with count below this cut off was replaced with zero since the sequence could not be differentiated from the noise (Galan *et al.*, 2018).

LFN_pos_: Based on relative frequency of sequences in positive control replicates with mock prey community, prey sequence that was mapped least frequently was used as a lower threshold *(Corse et al. 2017)*. Sequences in the dataset that were present below this threshold count were removed. We also got an unexpected sequence with taxonomic identification as leopard, which was not added to our mock prey community as the positive control. Hence, any sequence in the dataset that was identified as leopard was also discarded.

Table S2. Number of sequence reads retained during each step during HTS filtering.

|  | Total |
| --- | --- |
| Initial Reads | 15499038 |
| Obiclean | 15125149 |
| ID > 0.95 | 9694449 |
| Count > 10 | 9378898 |
| LFN_neg | 9310550 |
| LFN_pos | 9179632 |
| Final Count | 9060458 |

Table S3. Simper dissimilarity index to determine the contribution of different prey species in observed discrepancies in prey detections using metabarcoding and mechanical sorting approaches.

| **Species** | **average** | **sd** | **ratio** | **ava** | **avb** | **cumsum** | **p** |
| --- | --- | --- | --- | --- | --- | --- | --- |
| Chital | 0.154 | 0.139 | 1.107 | 0.686 | 0.237 | 0.253 | 0.001 ^***^ |
| Cattle | 0.15 | 0.153 | 0.981 | 0.763 | 0.417 | 0.5 | 0.001 ^***^ |
| Sambar | 0.136 | 0.15 | 0.904 | 0.641 | 0.506 | 0.724 | 1 |
| Nilgai | 0.058 | 0.11 | 0.528 | 0.205 | 0.071 | 0.82 | 0.012 ^*^ |
| Buffalo | 0.037 | 0.086 | 0.432 | 0.135 | 0.051 | 0.88 | 0.001 ^***^ |
| Wildpig | 0.026 | 0.074 | 0.347 | 0.096 | 0.026 | 0.923 | 0.004 ^**^ |
| Chinkara | 0.02 | 0.058 | 0.352 | 0.109 | 0.006 | 0.956 | 0.001 ^***^ |
| Sloth bear | 0.012 | 0.058 | 0.198 | 0.045 | 0 | 0.975 | 0.001 ^***^ |
| Primate | 0.009 | 0.04 | 0.239 | 0.051 | 0.006 | 0.991 | 0.001 ^***^ |
| Rodent | 0.003 | 0.02 | 0.139 | 0.019 | 0 | 0.996 | 0.001 ^***^ |
| Jungle fowl | 0.003 | 0.024 | 0.113 | 0.013 | 0 | 1 | 0.001 ^***^ |

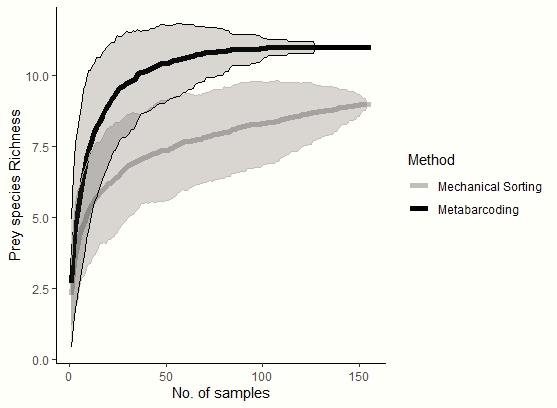

Figure S1. Sample-based rarefaction curve of the number of identified prey species in the diet of tigers in RTR.
